# Epilepsy protein myoclonin1 interacts with inositol 1,4,5–trisphosphate (IP_3_) receptor and reduces Ca^2+^ store in endoplasmic reticulum

**DOI:** 10.1101/2024.07.01.601633

**Authors:** Toshimitsu Suzuki, Kripamoy Aguan, Hideaki Mizuno, Takeshi Nakamura, Ikuyo Inoue, Katsuhiko Mikoshiba, Atsushi Miyawaki, Kazuhiro Yamakawa

## Abstract

Mutations of *EFHC1* gene have been identified in patients with epilepsies including juvenile myoclonic epilepsy (JME), and mice with *Efhc1* deficiency exhibit epileptic phenotypes. Myoclonin1 protein encoded by *EFHC1* is not expressed in neurons but in cells with motile cilia including those of choroid plexus and ependymal cells which form an epithelial layer lining brain ventricles. Detailed molecular basis of epilepsies caused by *EFHC1* mutations, however, remain unclear. Here we report that myoclonin1 is well co–expressed with inositol 1,4,5–trisphosphate receptor type 1 (IP_3_R1) at choroid plexus and ependymal cells and these two proteins bind each other. Endoplasmic reticulum (ER) of *Efhc1*–deficient mouse (*Efhc1*^-/–^) cells contains larger levels of calcium ions (Ca^2+^) than that of wild–type (WT) mice, and IP_3_–induced Ca^2+^ release (IICR) from ER is higher in *Efhc1*^-/–^ cells than that of WT. Furthermore, myoclonin1 revealed to interact with PRKCSH, also known as a protein kinase C substrate 80K–H which interacts with IP_3_R1. Myoclonin1 further binds to IP_3_R2 and IP_3_R3. Thus, our results indicate that myoclonin1 modulates ER–Ca^2+^ homeostasis through interactions with IP_3_Rs and PRKCSH, and suggest that myoclonin1 dysfunctions cause impaired intracellular Ca^2+^ mobilization. It’s relevance to the epileptic phenotypes of patients with *EFHC1* mutations is now of interest.

## Introduction

Human *EFHC1* gene encodes a 640 amino acid protein myoclonin1 harboring three tandemly repeated DM10 domains and one EF–hand calcium–binding motif at C terminus (Suzuki et al., 2004). We originally and mistakenly reported that myoclonin1 is expressed in neurons of mouse brain (Suzuki et al., 2004), but our subsequent studies using *Efhc1*^-/–^ mouse and a newly–developed mouse monoclonal antibody for myoclonin1 (6A3–mAb) revealed that myoclonin1 is actually not expressed in neurons but dominantly expressed in choroid plexus at embryonic stages, and at motile cilia of ependymal cells, tracheal motile cilia, and sperm flagella at postnatal stages (Suzuki et al., 2008; Suzuki et al., 2009; Suzuki et al., 2020). *EFHC1* heterozygous missense mutations in patients with epilepsies including JME have been repeatedly reported by us and many other groups (Suzuki et al., 2004; Ma et al., 2006; Stogmann et al., 2006; Annesi et al., 2007; Medina et al., 2008; Jara–Prado et al., 2012; Coll et al., 2016; Bailey et al., 2017; Raju et al., 2017; Thounaojam et al., 2017; Wang et al., 2017; Lin et al., 2023). A missense mutation in *EFHC1* has also been identified homozygously in patients with intractable epilepsy of infancy (Berger et al., 2012). We further reported that *Efhc1*– deficient mice exhibited spontaneous myoclonic seizures, increased seizure susceptibility to chemo– convulsant pentylenetetrazol (PTZ), decreased beating frequency of ependymal cells’ cilia as well as that of choroid plexus epithelial one and enlarged brain ventricles (Suzuki et al., 2009; Narita et al., 2012).

We have shown that over–expression of myoclonin1 in mouse hippocampal primary culture neurons activates R–type voltage–dependent calcium channel (Ca_v_2.3) resulting excessive intracellular Ca^2+^ influx and rapid cell–death, and these effects were compromised by *EFHC1*– mutations found in JME patients (Suzuki et al., 2004). We further reported that myoclonin1 interacts with transient receptor potential M2 channel (TRPM2) which is Ca^2+^–permeable cation channel and potentiates the channel activity (Katano et al., 2012). It is of interest whether myoclonin1 interacts with any other proteins to modulate Ca^2+^ signaling and homeostasis.

ER–Ca^2+^ homeostasis is maintained by IP_3_Rs and IP_3_, which induce release of Ca^2+^ from ER (Mikoshiba et al., 2007). There are three subtypes of IP_3_Rs, IP_3_R1–3, in mammals with distinct regulation and distribution throughout the body. IP_3_R1 is a predominant subtype in brain, and is involved in diverse functions such as development, axon–guidance and cognition (Ashworth et al., 2007; Xiang et al., 2002; Nagase et al., 2003; Tang et al., 2003). Because of the involvement of IP_3_R1 in epileptic phenotypes (Matsumoto et al., 1996) and the involvement of myoclonin1 in Ca^2+^ signaling as also suggested by its EF–hand Ca^2+^ binding motif (Suzuki et al., 2004), we investigated possible functional interplay between myoclonin1 and IP_3_R1. Here we report that in mouse brain, IP_3_R1 is well co–expressed with myoclonin1 at choroid plexus and ependymal cells and IP_3_R1 binds to myoclonin1. Myoclonin1–deficiency significantly increased levels of ER Ca^2+^ store ([Ca^2+^]_ER_) and IICR activity in cells from *Efhc1*^-/–^ mice. In addition, we find that myoclonin1 binds to PRKCSH which has been known to interact with IP_3_R1 (Kawaai et al., 2009). Our findings indicate that myoclonin1 regulates ER–Ca^2+^ homeostasis through interaction with IP_3_R1.

## Results

### Myoclonin1 and IP_3_R1 are co–expressed in choroid plexus and ependymal cells in brain

Immunohistochemical analyses revealed that myoclonin1 and IP_3_R1 were abundantly expressed in choroid plexus at embryonic day 14 (E14; **Fig. 1A–C**). As reported previously (Suzuki et al., 2008), the expression of myoclonin1 in choroid plexus was very transient, appearing at embryonic stages (E14–18) and gradually switching off until postnatal day 15 (P15; **Fig. 1D**). In contrast, the IP_3_R1 expression in choroid plexus was observed at E14 (**Fig. 1B, C**) and postnatal period (**Fig. 1D**), suggesting its continuous expression through embryonic and postnatal stages. At P15, myoclonin1 was prominently expressed in motile cilia of ependymal cells but also moderately in cell body (**Fig. 1D**); the somatic expression was more prominent in 3rd ventricle as reported previously (Suzuki et al., 2008). The IP_3_R1 expression was observed mainly in somata and not at cilia of ependymal cells (**Fig. 1D**). Altogether these results indicate that myoclonin1 and IP_3_R1 proteins are co–expressed in embryonic choroid plexus and postnatal ependymal cells.

**Figure 1.**
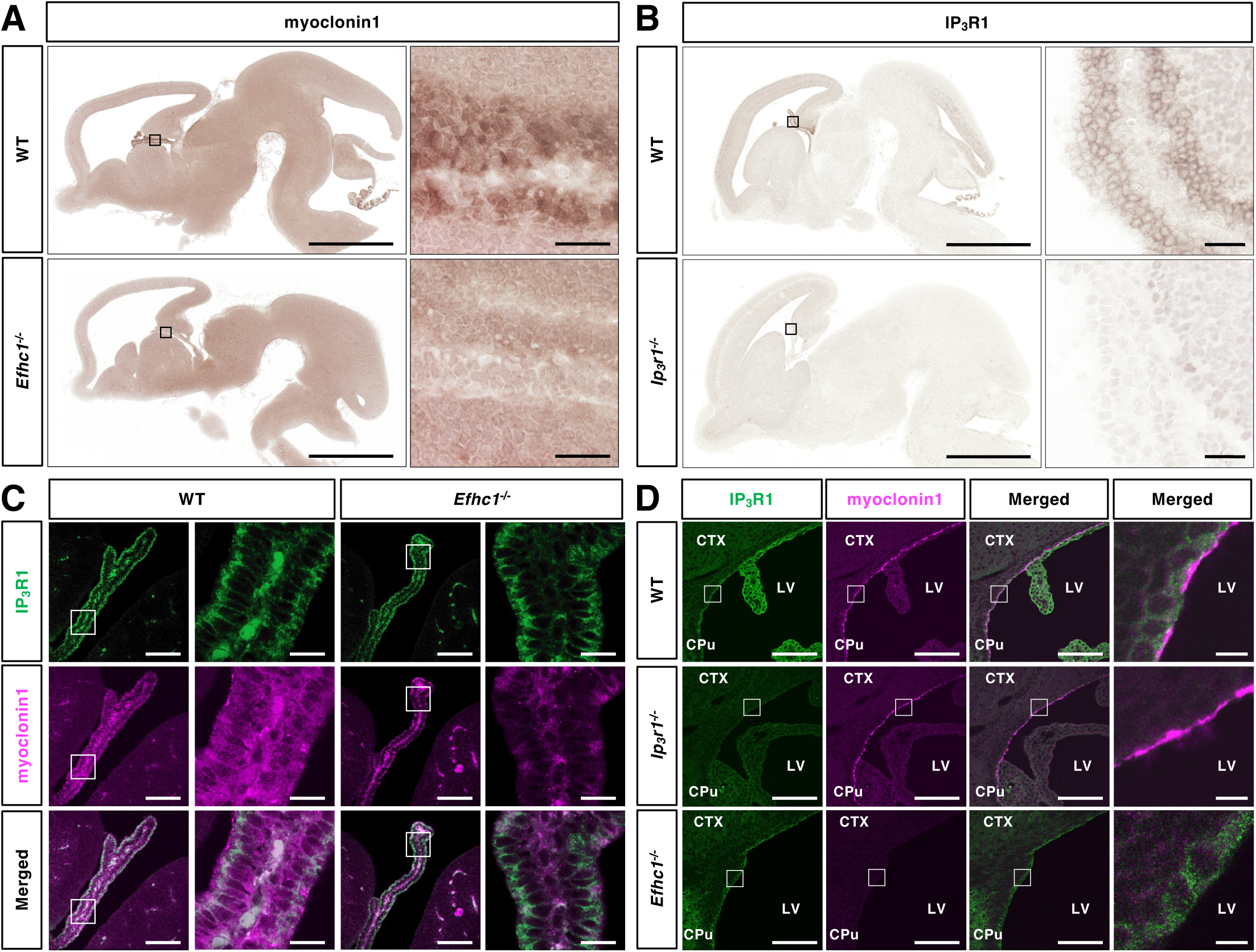
Co–expressions of myoclonin1 and IP_3_R1 in embryonic choroid plexus and postnatal ependymal cells. (**A–C**) Immunohistochemical analyses showed co–expression of myoclonin1 (brown in A, magenta in C) and IP_3_R1 (brown in B, green in C) in choroid plexus of mouse brain at E14 (sagittal brain sections, n=2 WT, 2 *Efhc1*^-/–^, 2 *Ip_3_r1*^-/–^). (**D**) Co–expressions of IP_3_R1 (green) and myoclonin1 (magenta) were observed in ependymal cells at P15 (sagittal sections, n=2 WT, 2 *Efhc1*^-/–^, 2 *Ip3r1*^-/–^). Boxed areas are enlarged. CTX, cortex; CPu, caudate putamen; LV, lateral ventricle. Scale bars= 1mm (low–magnification images in A, B), 25µm (high–magnification images in A, B), 100µm (low–magnification images in C, D), 15µm (high–magnification images in C) and 8µm (high–magnification images in D).

### Myoclonin1 interacts with all IP_3_R subtypes

We next investigated whether myoclonin1 binds to IP_3_Rs. Because a number of proteins interact with either of N– and C–terminal cytosolic regions of IP_3_R1 and modulate its activity (Mikoshiba et al., 2007), we selected these regions to test their ability to interact with myoclonin1 for co–immunoprecipitation (co–IP) assay. It actually revealed that myoclonin1 binds to C–terminal IP_3_R1 but not with N–terminal (**Fig. 2A, B**). By using a series of deletion constructs of the C– terminus, we further narrowed down the interacting region to amino acid residues (a.a.) 2565–2625 (**Fig. 2A, C**). Inversely, co–IP analyses using a series of deletion fragments of myoclonin1 revealed that each of the three DM10 domains of myoclonin1 independently bound to IP_3_R1 (**Fig. 2D, E**). C–terminal regions of three IP_3_R subtypes (IP_3_R1, IP_3_R2, IP_3_R3) are highly conserved (**Fig. S1A**), and a co–IP assay revealed that all C-termini of these IP_3_R subtypes similarly bound to myoclonin1 (**Fig. S1B**).

**Figure 2.**
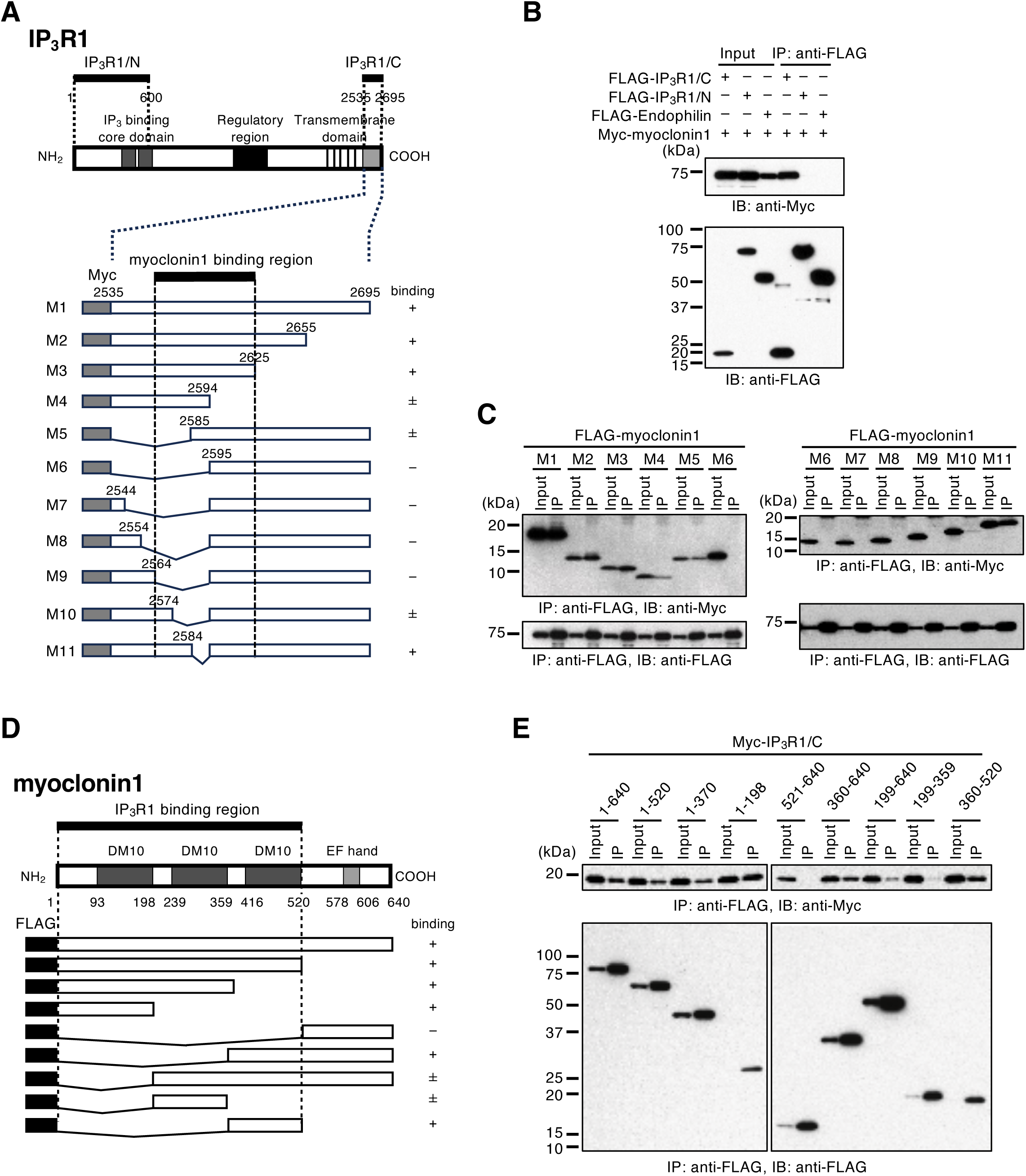
Myoclonin1 interacts with C-terminus of IP_3_R1. (**A**) Schematic diagrams of IP_3_R1, its deletion constructs and binding activity. Bold black lines indicate IP_3_R1 N– (a.a. 1–600) and C– (a.a. 2535–2695) terminus (top). A segment (a.a. 2565–2625; black line, middle) in C–terminal IP_3_R1 contains a binding site for myoclonin1. (**B**) Western blots of co–IP analysis showing that myoclonin1 interacted with IP_3_R1 C–terminus but not with N–terminus and Endophilin (negative control). (**C**) The interacting region of IP_3_R1 C–terminus to myoclonin1 was narrowed down to a.a. 2565–2625. (**D**) Schematic diagram of myoclonin1 deletion constructs and binding activity. A bold black line (top) contains a binding site for IP_3_R1. (**E**) Each of the three DM10 domains of myoclonin1independently bound to IP_3_R1. The degree of interaction is indicated by +, ±, –±, or – (A, D). /C, C–terminal; /N, N–terminal; IP, immunoprecipitation; IB, immunoblot; Input, 5% of cell lysate.

### Myoclonin1 interacts with an IP_3_R regulator PRKCSH

PRKCSH is a glucosidase 2 subunit beta which is also known as 80K-H. A previous report showed that PRKCSH was identified as an interacting protein of IP_3_Rs by yeast two-hybrid screening and regulates its activity (Kawaai et al., 2009). Because binding sites of IP_3_R1 for PRKCSH and myoclonin1 look overlapping, we investigated interactions among these three proteins. As previously reported, co–IP revealed that a.a. 2555–2594 of IP_3_R1 bound to PRKCSH (**Fig. S2**). A co-IP of myoclonin1 and a series of PRKCSH deletion constructs including two clones, a.a. 32–528 and 184–528 which were identified as fragments binding to IP_3_R1 (Kawaai et al., 2009) and additional newly designed one, revealed that a.a. 400–448 of PRKCSH binds to myoclonin1 (**Fig. 3A–E**). Similarly to IP_3_R1 (**Fig. 2D, E**), the three DM10 domains of myoclonin1 independently bound to PRKCSH (**Fig. 3F, G**).

**Figure 3.**
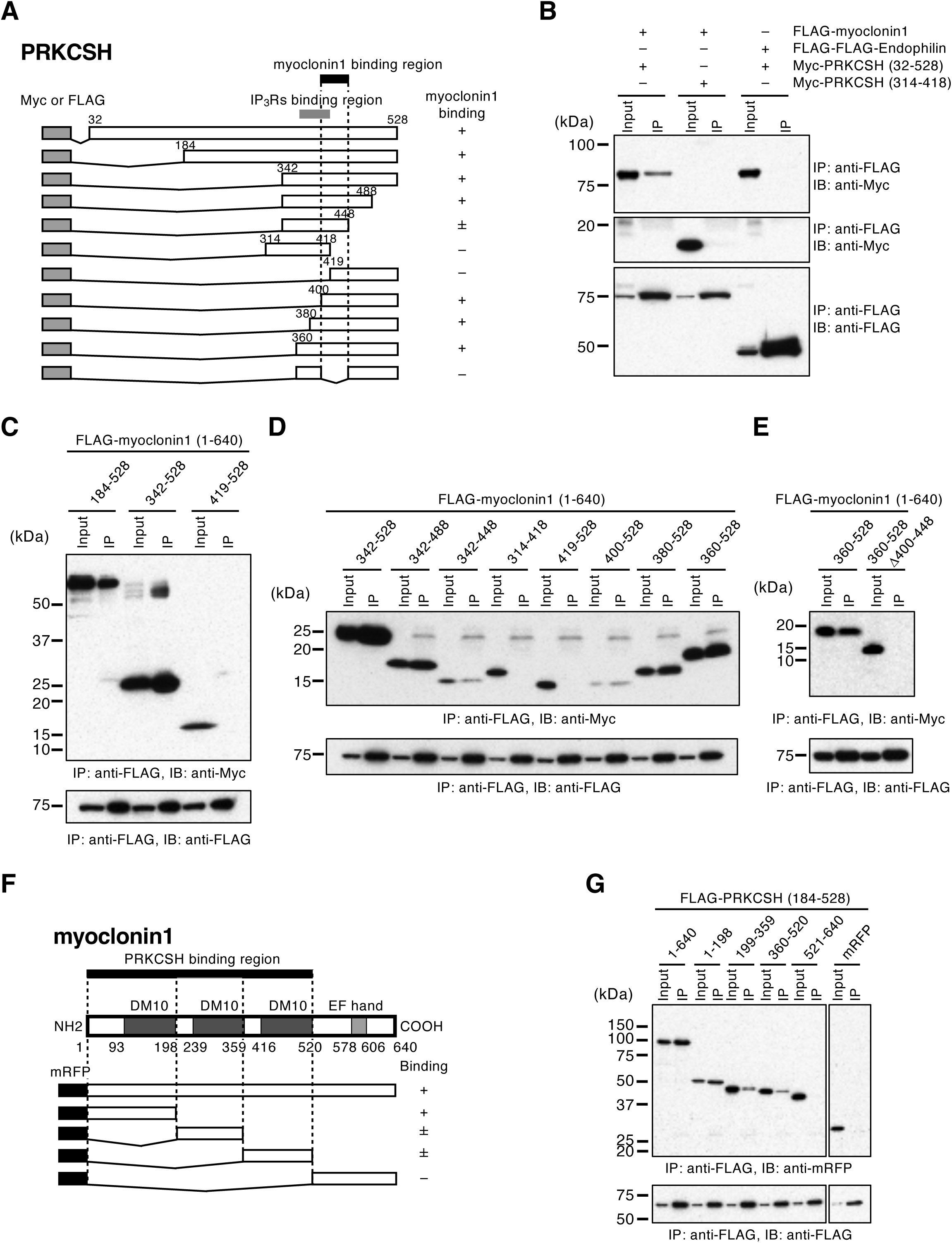
Myoclonin1 interacts with PRKCSH at its interaction site for IP_3_R1. (**A**) Schematic diagram of PRKCSH deletion constructs and binding activity. A short segment (a.a. 400–448; black line) in PRKCSH contains binding site for myoclonin1. Other one (a.a. 365–418; gray line) has been reported as a binding site for IP_3_Rs (Kawaai et al., 2009). (**B–E**) A region of PRKCSH a.a. 400–448 is critical to bind to myoclonin1. (**F**) Schematic diagram of myoclonin1 deletion constructs and binding activity. A bold black line (top) contains a binding site for PRKCSH. (**G**) PRKCSH bound to myoclonoin1 similarly to IP_3_R1. The degree of interaction is indicated by +, ± or – (A, F). IP, immunoprecipitation; IB, immunoblot; Input, 5% of cell lysate; Δ, internal deletion.

### Myoclonin1 regulates ER–Ca^2+^ store

We subsequently measured levels of [Ca^2+^]_ER_ in mouse embryonic fibroblasts (MEFs) and glial cells derived from *Efhc1*^-/–^ and WT littermates by applying ionomycin, an ionophore that induces formation of Ca^2+^–permeable pores in cellular membranes, leading to complete emptying of [Ca^2+^]_ER_ independently of IP_3_Rs activation (Singaravelu and Deitmer, 2006). The assay showed that [Ca^2+^]_ER_ levels of the *Efhc1*^-/–^ cells were significantly higher compared to WT (**Fig. 4A, B**). We also measured IICR by addition of bradykinin (BK), which stimulates phospholipase C (PLC) metabolism of phosphatidylinositol-4,5-biphosphate (PIPP) to IP_3_ (Lambert et al., 1986), and observed higher IICR level in *Efhc1*^-/–^ MEFs than in WT (**Fig. 4C**). Western blot analyses revealed that myoclonin1 expression was abrogated in the *Efhc1*^-/–^ MEFs, while those of IP_3_Rs and PRKCSH were remained unchanged (**Fig. S3**). These results indicate that myoclonin1 deficiency enhances [Ca^2+^]_ER_ and IICR.

**Figure 4.**
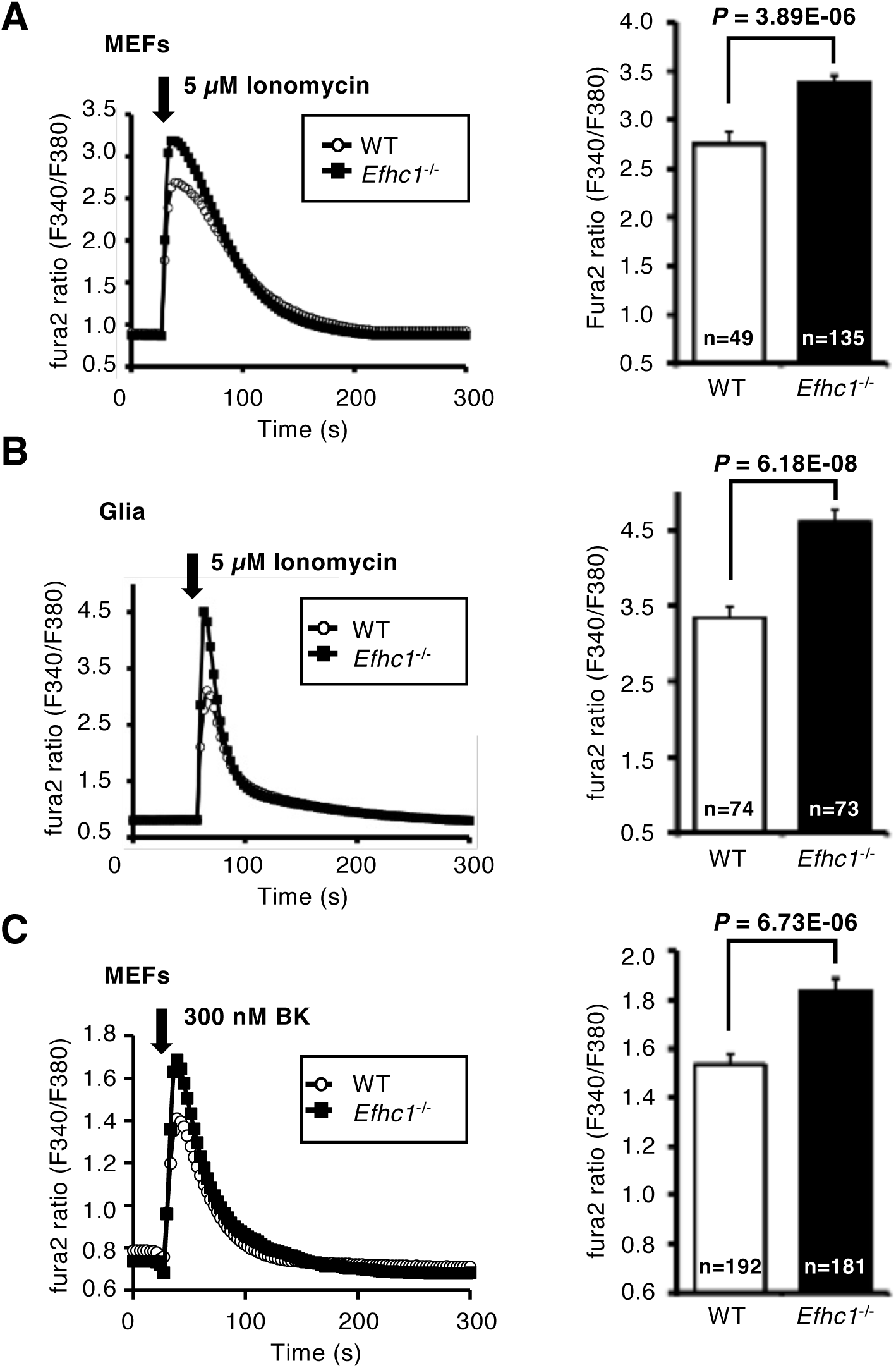
Myoclonin1 deficiency enhances [Ca^2+^]_ER_ and IICR. Ionomycin releasable [Ca^2+^]_ER_ was significantly higher in *Efhc1*^-/–^ cells than WT (**A**, n= 49 WT, 135 *Efhc1*^-/–^ MEFs; **B**, n= 74 WT, 73 *Efhc1*^-/–^ glial cells). (**C**) IICR induced by bradykinin (BK) was significantly higher in *Efhc1*^-/–^ MEFs than WT (n= 192 WT, 181 *Efhc1*^-/–^). Arrows indicate time point of addition of ionomycin or BK. n, total number of cells measured.

We further investigated whether myoclonin1 reduces [Ca^2+^]_ER_ and IICR. Human HeLa.S3 cells were transfected with monomeric red fluorescence protein (mRFP)–fused myoclonin1 which did not affect expressions of IP_3_Rs and PRKCSH (**Fig. S4A**). Ca^2+^ imaging revealed that [Ca^2+^]_ER_ by applying ionomycin and IICR by histamine, which generates IP_3_ through activation of PLC (Ishida et al., 2014), were significantly decreased by myoclonin1 over–expression (**Fig. S4B, C**). To confirm the effect, myoclonin1 was further re–introduced into *Efhc1*^-/–^ MEFs. Ca^2+^ imaging also revealed that both [Ca^2+^]_ER_ by ionomycin and IICR by BK were significantly attenuated in mRFP– myoclonin1 expressing MEFs compared to control one (**Fig. S4D, E**). Together, these results indicate that myoclonin1 lowers [Ca^2+^]_ER_ and is involved in maintenance of ER–Ca^2+^ homeostasis.

## Discussion

Here in this study, we showed that myoclonin1 forms a protein complex with IP_3_Rs and PRKCSH, and regulates IP_3_-mediated Ca^2+^ release. Together with our previous observations of the bindings of myoclonin1 with Ca_v_2.3 (Suzuki et al., 2004) and with TRPM2 (Katano et al., 2012), the results suggest that myoclonin1 is involved in intracellular Ca^2+^ mobilization.

A notable observation is that myoclonin1 and IP_3_R1 are co–expressed in ciliated cells such as choroid plexus and ependymal cells in the brain. We also found that myoclonin1 deficiency enhances [Ca^2+^]_ER_ and IICR in cells from *Efhc1*–deficient mice. It has previously been reported that, (1) Ca^2+^ initiates beating of cilia and flagella (Sataric et al., 2019), and (2) elevation of cytosolic Ca^2+^ contributes increased CBF of ependymal cells (Di Benedetto et al., 1991; Nguyen et al., 2001). Moreover, blockage of cilia–mediated cerebrospinal fluid (CSF) inflow has been known to be sufficient to predispose to seizures in another mouse model (Faubel et al., 2022). These may let us assume that reduced CBF by decreased cytosolic Ca^2+^ causes seizures in the *Efhc1*–deficient mice. However, in our previous study of the mice we showed that reduced CBF was observed only in homozygous *Efhc1*^-/–^ mice but not in heterozygous one (*Efhc1*^+/–^) which still showed seizure phenotypes such as frequent spontaneous myoclonus and increased seizure susceptibility to PTZ (Suzuki et al., 2009). These results indicate that there is considerable inconsistency between seizure phenotypes and CBF in heterozygous *Efhc1*^+/–^ mice. Taken together, these observations may suggest that blockage of cilia–mediated CSF inflow and reduction of CBF themselves may not or minimally contribute directly to the seizure phenotypes observed in the *Efhc1*–deficient mice, and an alternative pathway would be required to explain the molecular basis of epilepsies for the mice.

We have reported that myoclonin1 is dominantly expressed in choroid plexus epithelial cells at embryonic stage (Suzuki et al., 2008), and in this study we found that myoclonin1 and IP_3_R1 are well co–expressed in those cells as well and these two proteins bind each other as mentioned above. In addition, we also have reported that CBF of neonatal choroid plexus epithelial cells from *Efhc1*– deficient mice was significantly lower than that of WT mice (Narita et al., 2012). Based on these findings we assume that myoclonin1 possibly plays critical role in choroid plexus epithelial cells. Further, the cells synthesize neurotrophic factors and other signaling molecules including insulin that are secreted in response to increased intracellular Ca^2+^ level (Mazucanti et al., 2019). Together with previous observation that IP_3_R1–deficient mice suffer from epilepsy (Matsumoto et al., 1996) and our results of Ca^2+^ imaging analyses which revealed increased [Ca^2+^]_ER_ and IICR through IP_3_R1 function in cells from *Efhc1*–deficient mice, it is likely that impaired IP_3_R1–mediated Ca^2+^ signaling in the mice affects secretion of several signaling molecules from choroid plexus cells leading to the altered neural circuit formation that might be responsible for epileptic phenotypes.

In addition to *EFHC1*, we have identified and reported multiple pathogenic mutations of *CILK1* (*ciliogenesis associated kinase 1*) gene, formerly known as *ICK* (*intestinal–cell kinase*), in patients affected with JME from a number of families (Bailey et al., 2018). High level of CILK1 expression was observed in ependymal and choroid plexus cells in postnatal mouse brain (Bailey et al., 2018; Tsutsumi et al., 2018). Moreover, pathogenic mutations of several another ciliogenesis associated proteins, CDKL5/STK9 (Kalscheuer et al., 2003; Tao et al., 2004), SDCCAG8 (Otto et al., 2010; Yamamura et al., 2017), PRICKLE1 and PRICKLE2 (Bassuk et al., 2008; Tao et al., 2011) have also been reported in patients with epilepsies including JME. These observations support the notion that functional impairments of the cells with motile cilia in brain, such as cell–cell communication, synthesis of neurotrophic factors and signaling molecules, secretion of the molecules, and CSF or ion homeostasis, are likely the basis of epileptic seizure phenotypes in patients with *EFHC1* mutations.

Our results presented here indicate that myoclonin1 participates in controlling intracellular Ca^2+^ mobilization in a manner that involves IP_3_Rs. This could be molecular basis underlying pathology of epilepsies caused by *EFHC1–*mutations and potential candidate for prevention strategies and treatments of epilepsies.

## Materials and Methods

### Study approval

All animal experimental protocols were approved by the Animal Experiment Committee of RIKEN and by the Institutional Animal Care and Use Committee of the Nagoya City University (NCU). All animal breeding and experimental procedures were performed in accordance with the ARRIVE guidelines and regulations of the RIKEN and the NCU.

### Mice

*Efhc1*– and *Ip_3_r1*–deficient mice used for this study have been developed and reported previously (Suzuki et al. 2009; Matsumoto et al., 1996). The heterozygous mice were maintained on C57BL/6J background, and resultant heterozygous mice were interbred to obtain WT, heterozygous, and homozygous mice.

### Ca^2+^ imaging

Ca^2+^ imaging analyses were done as described previously (Cai et al., 2004). We placed the cells on an inverted microscope IX70 (Olympus) and observed through an objective lens UApoN40XO340, etc. (Olympus). Some of images were acquired using Olympus IX81–ZDC. The cells were illuminated by a Xenon lamp through excitation filters, 340AF15 (Omega) and 380HT15 (Omega), alternately. We used a computer-controlled filter exchanger Lambda 10-2 (Sutter) to switch filters. We used a dichroic mirror and an emission filter 430DCLP (Omega) and 510WB40 (Omega), respectively. To eliminate Ca^2+^ influx, we performed all experiments in absence of extracellular Ca^2+^. We measured IICR with addition of 5µM histamine (Ishida et al., 2014) for HeLa.S3 cells or 300nM BK (Lambert et al., 1986) for MEFs whereas ER–stored Ca^2+^ was measured with addition of 5µM ionomycin (Singaravelu and Deitmer, 2006). All measurements shown were representative results from two ∼ four independent experiments (∼3 dishes per condition were used in each experiment).

### Statistical analysis

Data were presented as mean ± standard error of the mean (s.e.m.) and statistical significance of differences between means was tested using unpaired t–test or analysis of variance (ANOVA) for repeated measures, using Fisher’s protected least–significance (PLSD) test. Statistical significance levels were defined as P < 0.05.

**Details of the other experiments were described in the Supplemental data.**

## CRediT authorship contribution statement

**Toshimitsu Suzuki:** Conceptualization, Investigation, Writing – original draft, Visualization, Funding acquisition. **Kripamoy Aguan:** Conceptualization, Investigation, Writing – original draft, Visualization. **Hideaki Mizuno:** Conceptualization, Methodology, Investigation, Resources, Writing – review & editing, Visualization. **Takeshi Nakamura:** Conceptualization, Investigation, Resources. **Ikuyo Inoue:** Investigation, Writing – review & editing. **Katsuhiko Mikoshiba:** Resources, Writing – review & editing, Funding acquisition. **Atsushi Miyawaki:** Resources, Writing – review & editing, Funding acquisition. **Kazuhiro Yamakawa:** Conceptualization, Resources, Writing – review & editing, Supervision, Funding acquisition.

## Declaration of competing interest

We have no competing interests to declare.

## Acknowledgements

We thank members of Laboratories for Cell Function Dynamics, Developmental Neurobiology, and Neurogenetics at RIKEN Center for Brain Science (CBS), Department of Neurodevelopmental Disorder Genetics and animal facility at the Nagoya City University (NCU). We are indebted to the Support Unit for Bio–material Analysis, Research Resources Division (RRD) at the RIKEN CBS and the Research Equipment Sharing Center at the NCU. We are also grateful to RIKEN CBS–Olympus Collaboration Center.

## Funding

This work was partly supported by grants from RIKEN CBS (K.M., A.M, and K.Y.), from NCU (K.Y.), from the JSPS KAKENHI (Grant Numbers 20790866 and 22791154 to T.S.) and from The Japan Epilepsy Research Foundation (T.S.).

## Notes

### Competing Interest Statement

The authors have declared no competing interest.

