## Supplementary data for "Epilepsy protein myoclonin1 interacts with inositol 1,4,5–trisphosphate (IP_3_) receptor and reduces Ca^2+^ store in endoplasmic reticulum"

**Supplemental Methods**

**Expression constructs.** *EFHC1* clone was described in previously [1]. We amplified parts of $IP_3R1$ (GenBank: NM_002222), $IP_3R2$ (NM_002223), $IP_3R3$ (NM_002224), PRKCSH (NM_002743) from human adult brain cDNA (Clontech) by PCR using KOD–plus (TOYOBO) and cloned them into pcDNA3–MycN, pcDNA3–FlagN or pcDNA3–mRFP vectors. We introduced mutations by using QuickChange Site–Directed Mutagenesis kit (Agilent Technologies) and confirmed nucleotide changes as well as integrity of full sequences by DNA sequencing.

**Antibodies**. Mouse monoclonal anti–myoclonin1 antibody (6A3–mAb) was reported previously [2–3]. Following antibodies were also used: $IP_3R1$ (KM1112 [4], 18A10 [5], 10A6 [6], H–80 or C–20, Santa Cruz Biotechnology), $IP_3R2$ (KM1083 [4]), $IP_3R3$ (KM1083 [4] and BD Transduction lab), FLAG (Sigma), Myc (Cell Signaling), GAPDH (Santa Cruz Biotechnology), PRKCSH (Santa Cruz Biotechnology) and DsRed (Invitrogen).

**Cell culture and transfection.** Mouse embryonic fibroblasts (MEFs) were prepared as described previously from E16 mouse tail [7]. Once the cells were confluent in poly–L–Lysine coated 60 mm dish (IWAKI), it was stored at –80 °C until needed. To obtain glial cells, culture medium of hippocampal neuron culture prepared from E16 mouse brain [1–2] was replaced with Dulbecco's Modified Eagle Medium (D–MEM) +10% fetal bovine serum (FBS) + 30 U/ml penicillin and 30 mg/ml streptomycin (P/S) at 2 days in vitro (DIV) and cultured for 18 DIV. The glial cells were harvested and stored in –80 °C for subsequent assay. MEFs and HeLa.S3 cells were transfected using Lipofectamine

LTX and PLUS reagent (Thermo Fisher Scientific) according to manufacturer protocol.

**Immunohistochemistry.** Preparations of mouse sagittal brain sections (paraffin) and immunohistochemical staining were carried out as described previously [2–3]. Antibodies, 18A10 for $IP_3R1$ and 6A3–mAb for myoclonin1, were used for staining. Images were acquired using AX80 (Olympus) or TCS SP2 (Leica) microscopes.

**Immunoprecipitation and western blot analysis.** We washed cultured cells twice in PBS and lysed in lysis buffer [10 mM Tris–HCl, pH 7.5, 150 mM NaCl, 1 mM EDTA, 0.5 or 1% NP40 and supplemented with protease inhibitor (Complete, Roche)]. Co–immunoprecipitation studies were carried out as described previously [1]. We probed blot with anti–FLAG, anti–Myc or anti–DsRed antibodies and developed with Western Lighting kit (Perkin Elmer). Wherever necessary, we stripped blots and re–probed with respective antibodies.

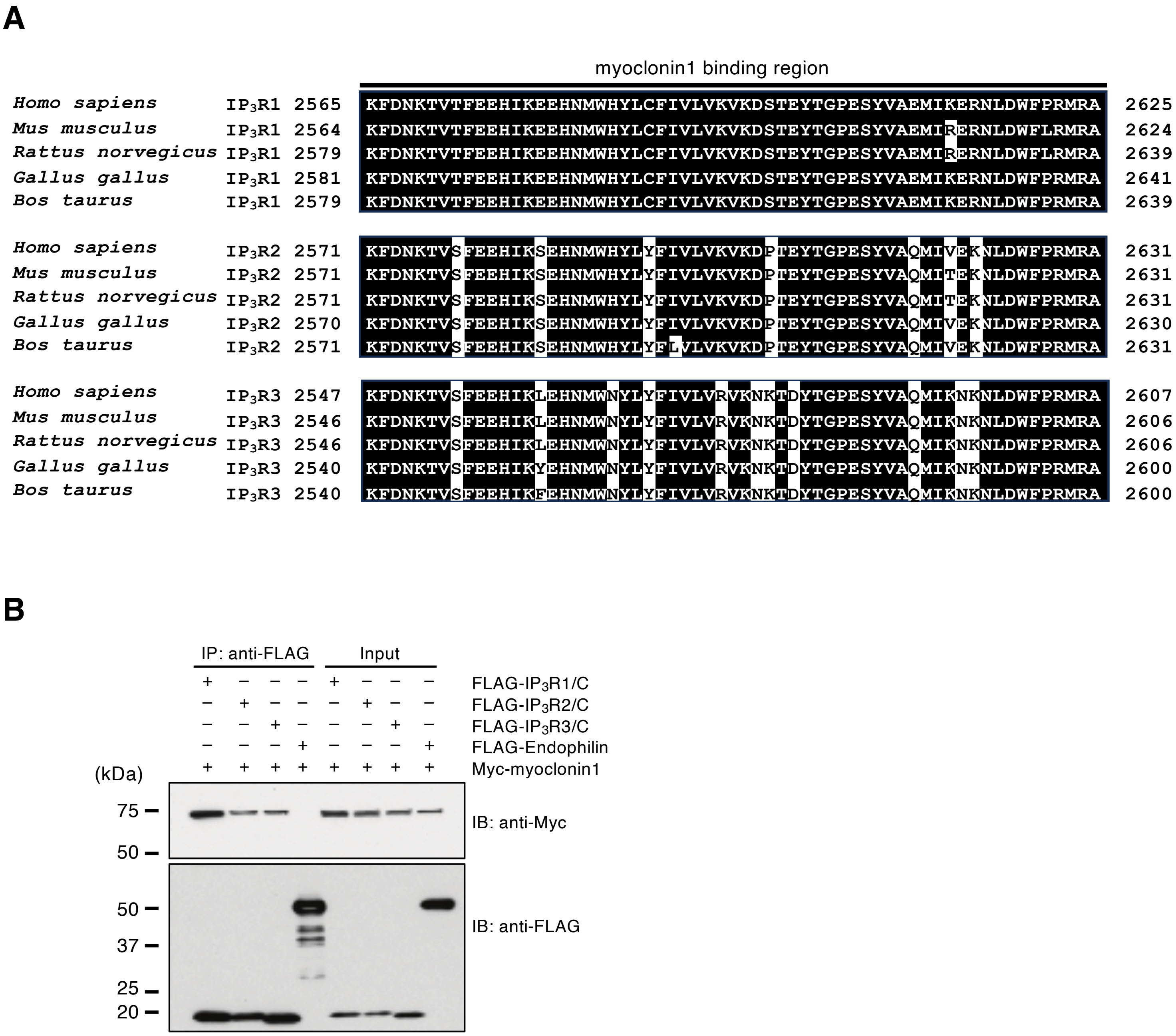


**Figure S1. Myoclonin1 interacts with C–termini of all three IP$_3$R subtypes.** (**A**) Binding regions of all three IP$_3$R subtypes (IP$_3$R1, 2, 3) to myoclonin1 are highly conserved. (**B**) IP$_3$R2 and IP$_3$R3 C-termini also interacted with myoclonin1 but not with Endophilin (negative control). IP, immunoprecipitation; IB, immunoblot; Input, 5% of cell lysate; /C, C–terminal.

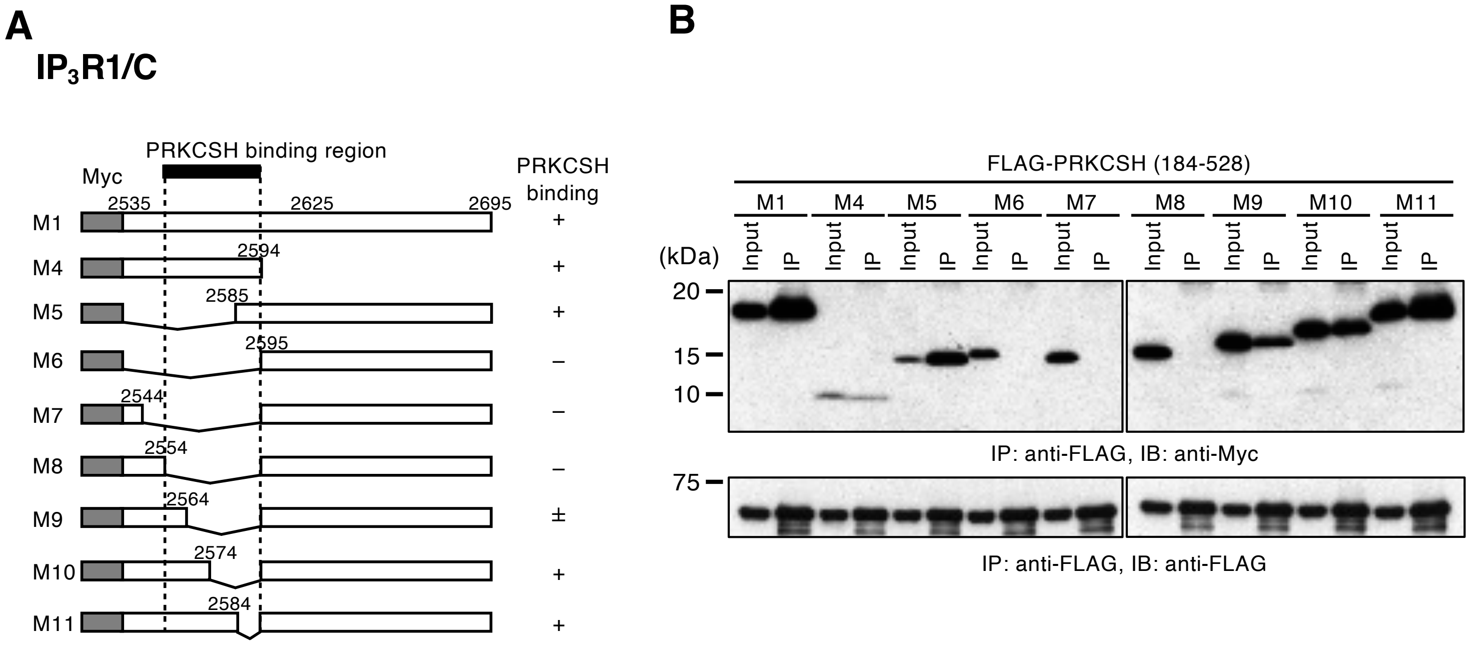


**Figure S2. PRKCSH interacts with IP$_3$R1 C-terminus at its interaction site for myoclonin1.** (**A**) Schematic diagram of C–terminal IP$_3$R1 deletion constructs and their binding activities to PRKCSH. A bold black bar (top) indicates a binding site to PRKCSH. Degrees of interaction are indicated by +, $\pm$ or –. (**B**) Western blots of co-IP showing that PRKCSH bound to a.a. 2555–2594 of C–terminal IP$_3$R1. IP, immunoprecipitation; IB, immunoblot; Input, 5% of cell lysate; /C, C–terminal.

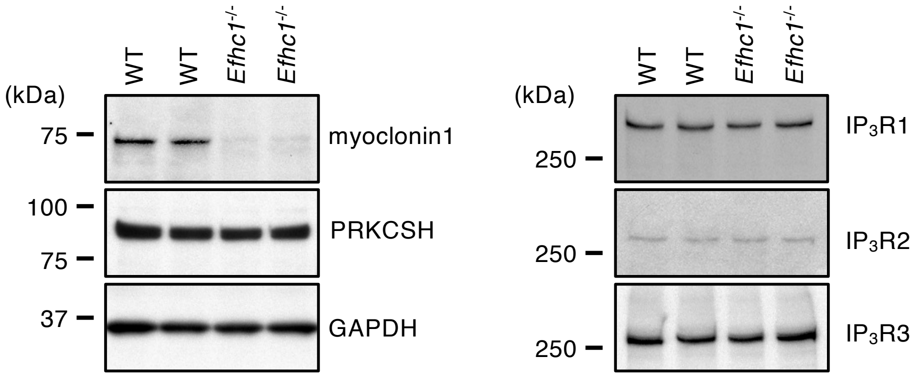


**Figure S3. Expressions of IP$_3$Rs and PRKCSH do not change in *Efhc1*$^{-/-}$ MEFs.** Western blot analyses revealed that myoclonin1 expression was abrogated in *Efhc1*$^{-/-}$ MEFs, whereas expressions of endogenous IP$_3$Rs and PRKCSH do not change (n=2 independent embryos per genotypes). An antibody to GAPDH was used as a control.

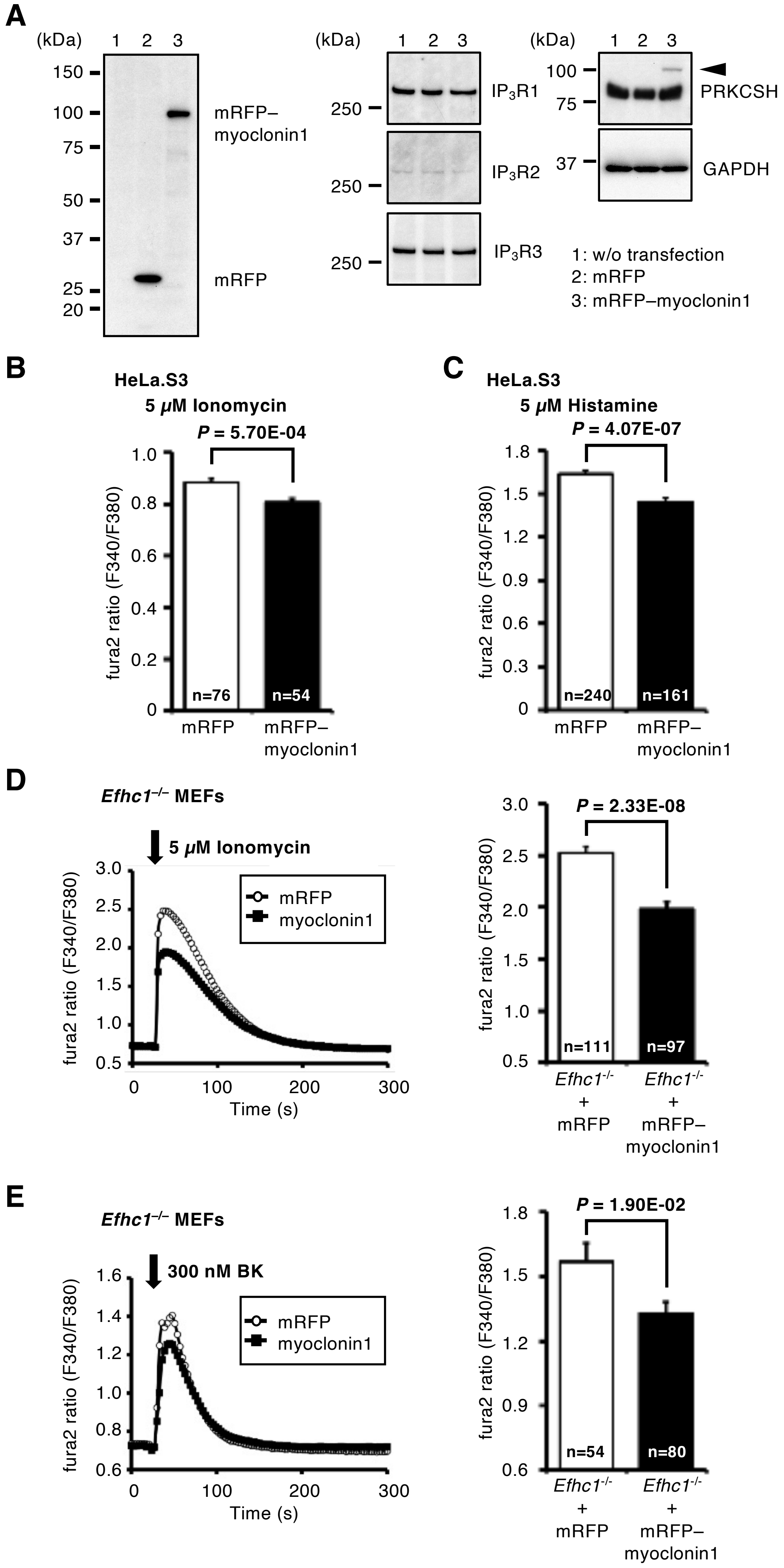
A
(kDa) 1 2 3
150
100
75
50
37
25
20
mRFP–
myoclonin1
mRFP
(kDa) 1 2 3
250
IP3R1
250
IP3R2
250
IP3R3
(kDa) 1 2 3
100
75
PRKCSH
37
GAPDH
1: w/o transfection
2: mRFP
3: mRFP–myoclonin1
B
HeLa.S3
5 μM Ionomycin
P = 5.70E-04
fura2 ratio (F340/F380)
n=76
n=54
mRFP
mRFP–
myoclonin1
C
HeLa.S3
5 μM Histamine
P = 4.07E-07
fura2 ratio (F340/F380)
n=240
n=161
mRFP
mRFP–
myoclonin1
D
Efhc1–/– MEFs
5 μM Ionomycin
fura2 ratio (F340/F380)
mRFP
myoclonin1
Time (s)
P = 2.33E-08
n=111
n=97
Efhc1-/- + mRFP
Efhc1-/- + mRFP–myoclonin1
E
Efhc1–/– MEFs
300 nM BK
fura2 ratio (F340/F380)
mRFP
myoclonin1
Time (s)
P = 1.90E-02
n=54
n=80
Efhc1-/- + mRFP
Efhc1-/- + mRFP–myoclonin1

**Figure S4. Myoclonin1 reduces $[Ca^{2+}]_{ER}$ and IICR.** (**A**) Western blot analyses [n=1 without (w/o) transfection, 1 mRFP, 1 mRFP–myoclonin1] revealed that exogenous of mRFP or mRFP–myoclonin1 proteins were detected at expected size (~30 kDa and ~100 kDa, respectively). Expression levels of endogenous $IP_3Rs$ and PRKCSH were not affected by over–expression of myoclonin1 in the cells. An mRFP–myoclonin1 band at ~100 kDa remained weakly in blot for PRKCSH (arrowhead) even after stripping of blot. An antibody to GAPDH was used as a control. (**B, C**) Ionomycin releasable $[Ca^{2+}]_{ER}$ (B; n=76 mRFP, 54 mRFP–myoclonin1 expressing cells) and histamine evoked IICR (C; n=240 mRFP, 161 mRFP–myoclonin1 expressing cells) were significantly lower in mRFP–myoclonin1 expressing HeLa.S3 cells than mRFP expressing one. Ionomycin releasable $[Ca^{2+}]_{ER}$ (**D**, n=111 mRFP, 97 mRFP–myoclonin1) and bradykinin (BK) evoked IICR (**E**, n=54 mRFP, 80 mRFP–myoclonin1) were significantly lower in mRFP–myoclonin1 expressing *Efhc1*$^{-/-}$ MEFs than mRFP expressing one. Arrows indicate the time point of addition of ionomycin or BK. n, total number of cells measured.
